## Supplementary material for "Perturbations in eIF3 subunit stoichiometry alter expression of ribosomal proteins and key components of the MAPK signaling pathways": all in one

### Supplementary Figures

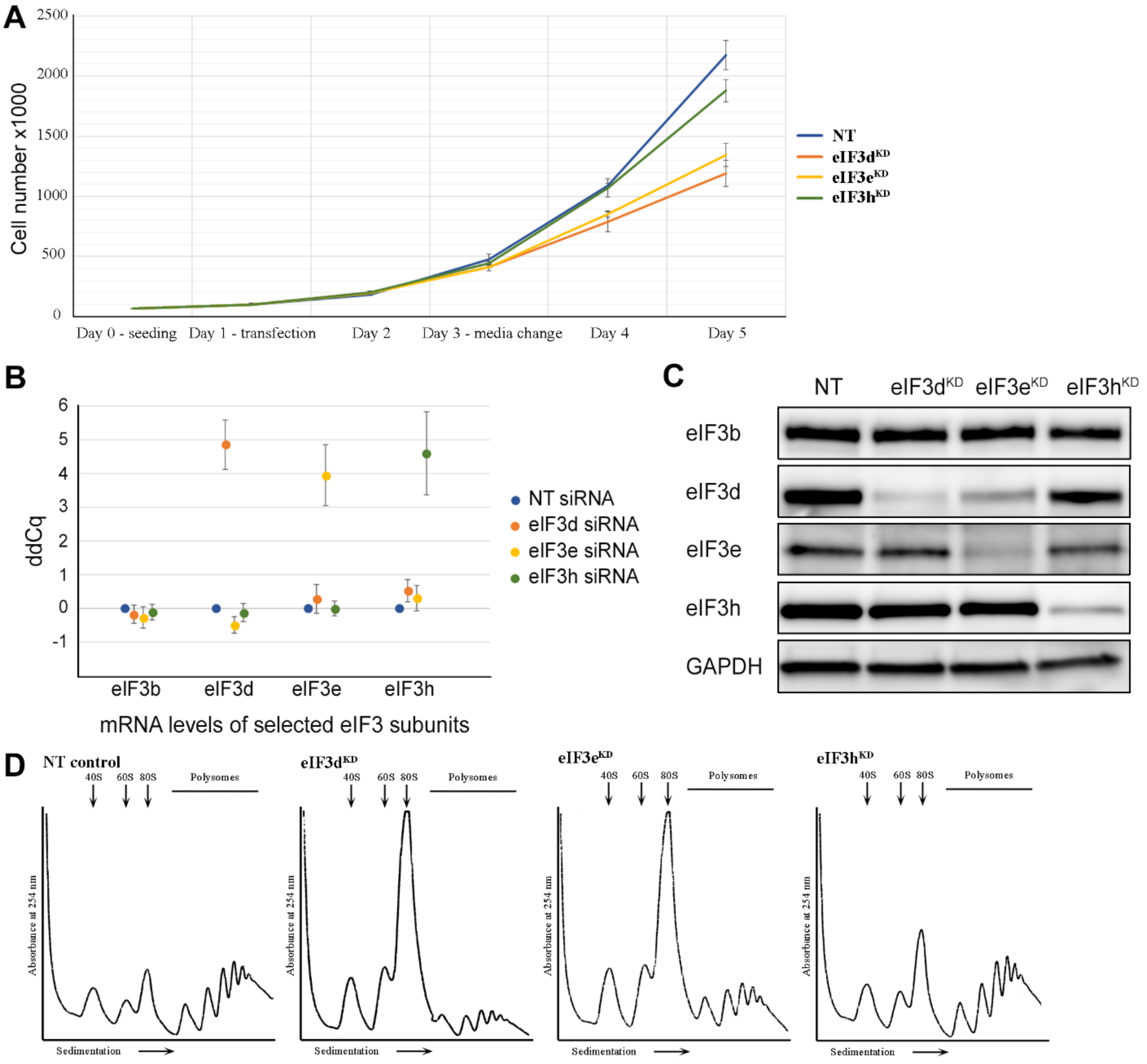


Supplementary Figure 1. **Quality control of eIF3e, d, and h knock-downs - Hela cells downregulated for eIF3d, eIF3e and eIF3h display slower growth and impaired translation. (A)** Growth curve of Hela cells transfected with siRNA targeting eIF3d, eIF3e, eIF3h or non-targeting (NT) siRNA as a control. Cells were grown in 6 well plates in duplicates and counted by Corning cell counter. **(B)** mRNA levels of the selected downregulated eIF3 subunits were significantly decreased as assessed by quantitative PCR for all replicates. Plot represents the result of all replicates used for RNA-Seq and Ribo-Seq library preparations ± SD. The ddCq value displays the threshold cycle normalized to the reference gene ALAS1 and to control cells transfected with non-targeting siRNA (NT). The ddCq values are in log2 scale. A ddCq of 0 indicates no change to NT siRNA control; ddCq = 4 indicates a drop to 6.25% (16 fold) compared to NT siRNA control. **(C)** Downregulation of selected eIF3 subunits targeted by siRNA was determined by Western blotting with GAPDH used as a loading control and antibodies listed in Supplementary Table S3. **(D)** Standard polysome profiles illustrate that cells were not overgrown at the time of harvest and occurred in a state of active translation (NT) with the expected specific reduction in polysomes in all three knock-downs, as reported previously (Wagner *et al*, 2016). One example profile for each knock-down is shown.

**A** **Spearman correlation between replicates**

| nt_FP | rep_1 | rep_2 |
|---|---|---|
| rep_1 | | |
| rep_2 | 0.989 | |
| rep_3 | 0.985 | 0.986 |

| d_FP | rep_1 | rep_2 |
|---|---|---|
| rep_1 | | |
| rep_2 | 0.981 | |
| rep_3 | 0.976 | 0.984 |

| e_FP | rep_1 | rep_2 |
|---|---|---|
| rep_1 | | |
| rep_2 | 0.984 | |
| rep_3 | 0.982 | 0.986 |

| h_FP | rep_1 | rep_2 |
|---|---|---|
| rep_1 | | |
| rep_2 | 0.986 | |
| rep_3 | 0.985 | 0.989 |

| nt_mRNA | rep_1 | rep_2 | rep_3 |
|---|---|---|---|
| rep_1 | | | |
| rep_2 | 0.994 | | |
| rep_3 | 0.988 | 0.990 | |
| rep_4 | 0.994 | 0.994 | 0.990 |

| d_mRNA | rep_1 | rep_2 | rep_3 |
|---|---|---|---|
| rep_1 | | | |
| rep_2 | 0.984 | | |
| rep_3 | 0.992 | 0.989 | |
| rep_4 | 0.992 | 0.986 | 0.993 |

| e_mRNA | rep_1 | rep_2 | rep_3 |
|---|---|---|---|
| rep_1 | | | |
| rep_2 | 0.990 | | |
| rep_3 | 0.993 | 0.993 | |
| rep_4 | 0.994 | 0.991 | 0.994 |

| h_mRNA | rep_1 | rep_2 | rep_3 |
|---|---|---|---|
| rep_1 | | | |
| rep_2 | 0.994 | | |
| rep_3 | 0.992 | 0.993 | |
| rep_4 | 0.989 | 0.988 | 0.988 |

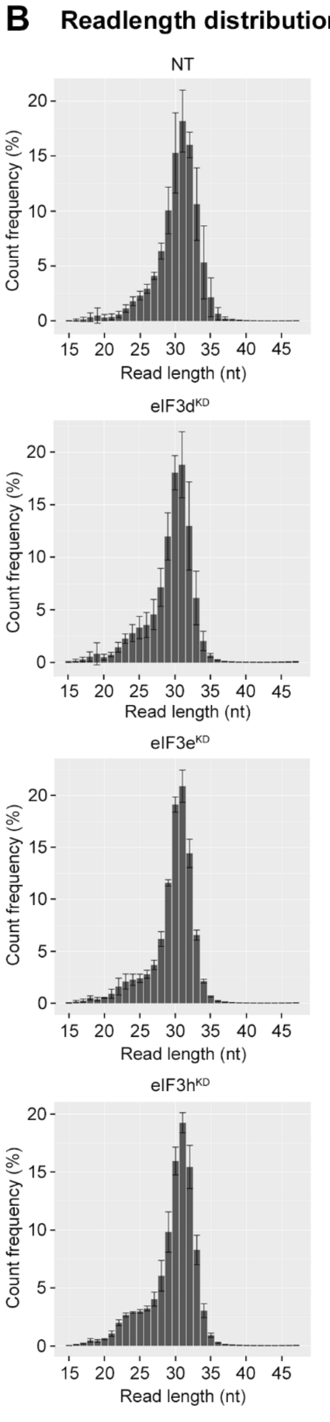


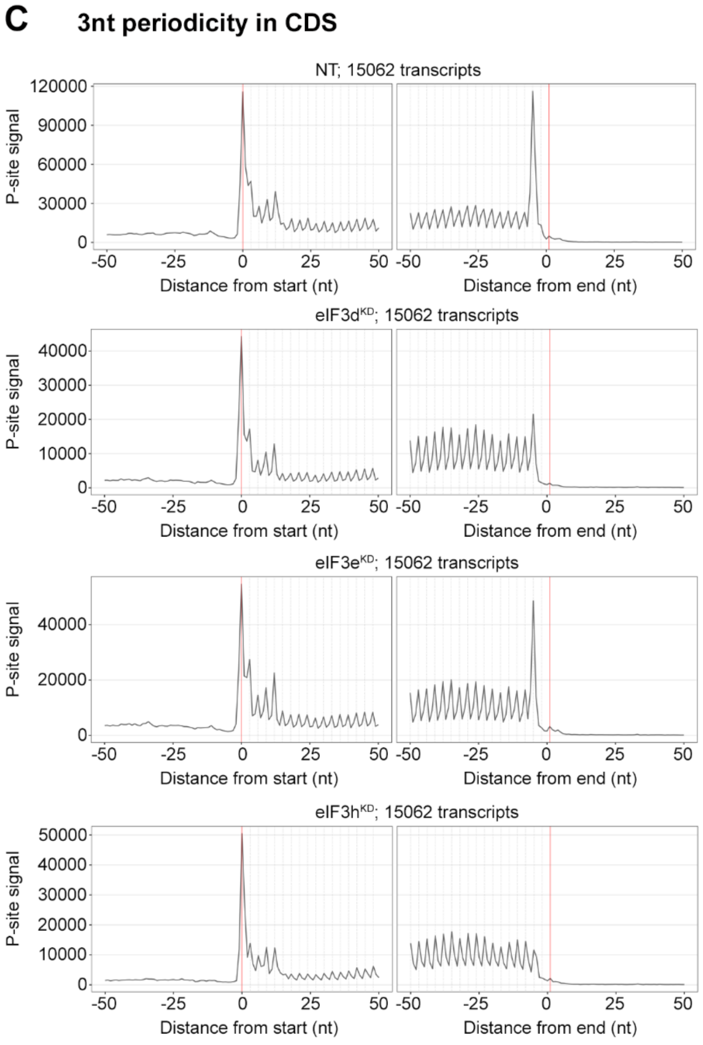


Supplementary Figure 2. **Quality control of Ribo-Seq libraries from all eIF3 knock-downs and the NT control. (A)** Spearman correlation coefficients of the footprint or mRNA count per gene among all replicates. **(B)** Read length distribution of footprints in Ribo-Seq libraries from all eIF3 knock-downs and the NT control. Bar charts show average values for each fragment length (after alignment to genome); all Ribo-Seq libraries were done in triplicates. **(C)** Meta-profiles showing the periodicity of ribosomes along the transcripts at the genome-wide scale. The metaprofiles are based on the P-site identification obtained by using riboWaltz (Lauria *et al*, 2018).

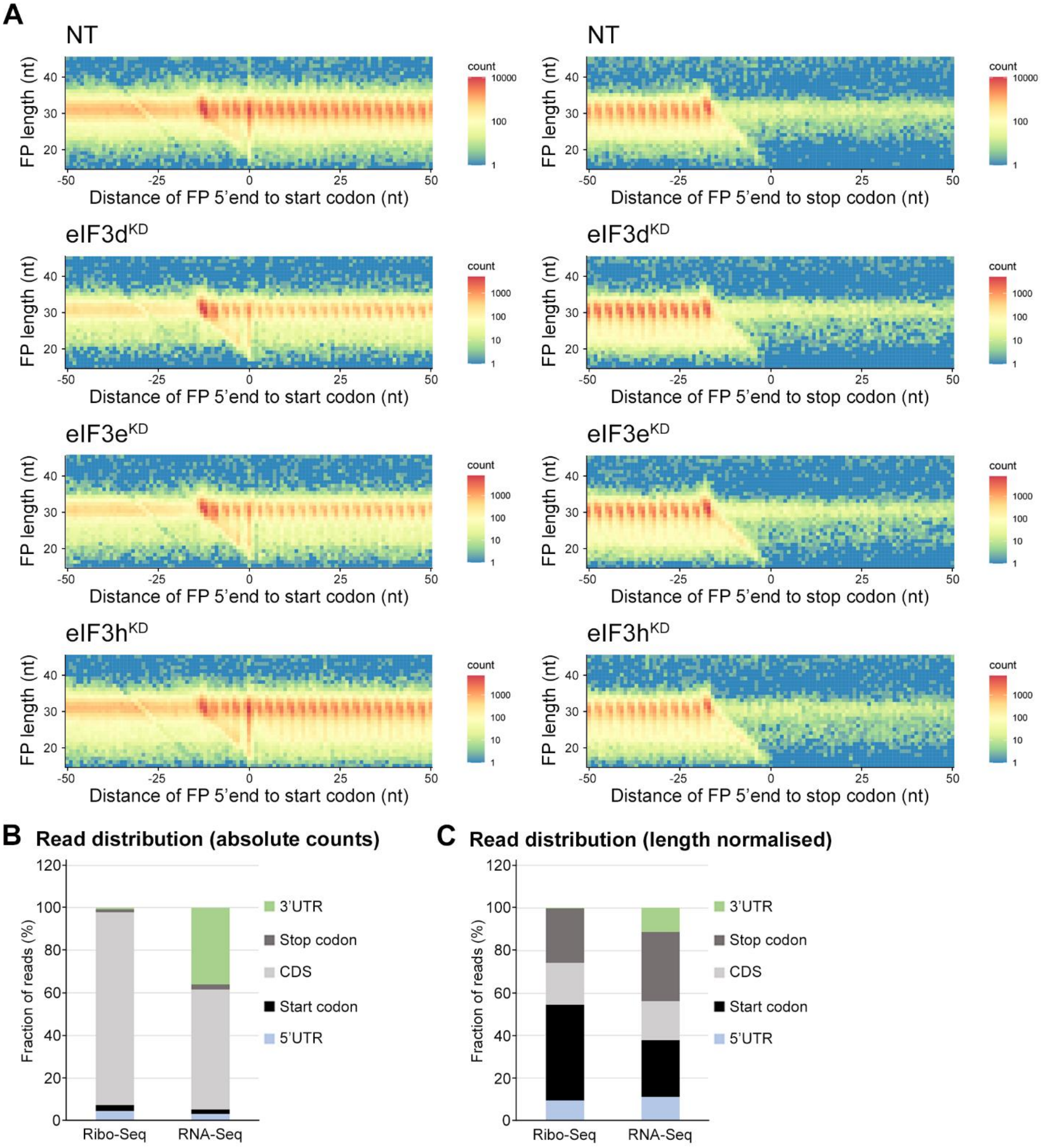


Supplementary Figure 3. **Ribo-Seq libraries from all eIF3 knock-downs and the NT control display triplet periodicity and enrichment of reads in CDS. (A)** All Ribo-Seq libraries show typical 3nt periodicity in the CDS. Metagene plots of footprint (FP) length versus 5' end position relative to the first nucleotide (position 0) of start (left) or stop codons (right). The color scale represents FP count as indicated on the right and is plotted in log scale. The labels at the color bar are given in linear scale. Heatmaps are shown for one exemplary replicate from each sample. **(B)** Fraction of reads in different transcript features **(C)** Same as in (B) but numbers of reads in each feature were normalized for average feature length.

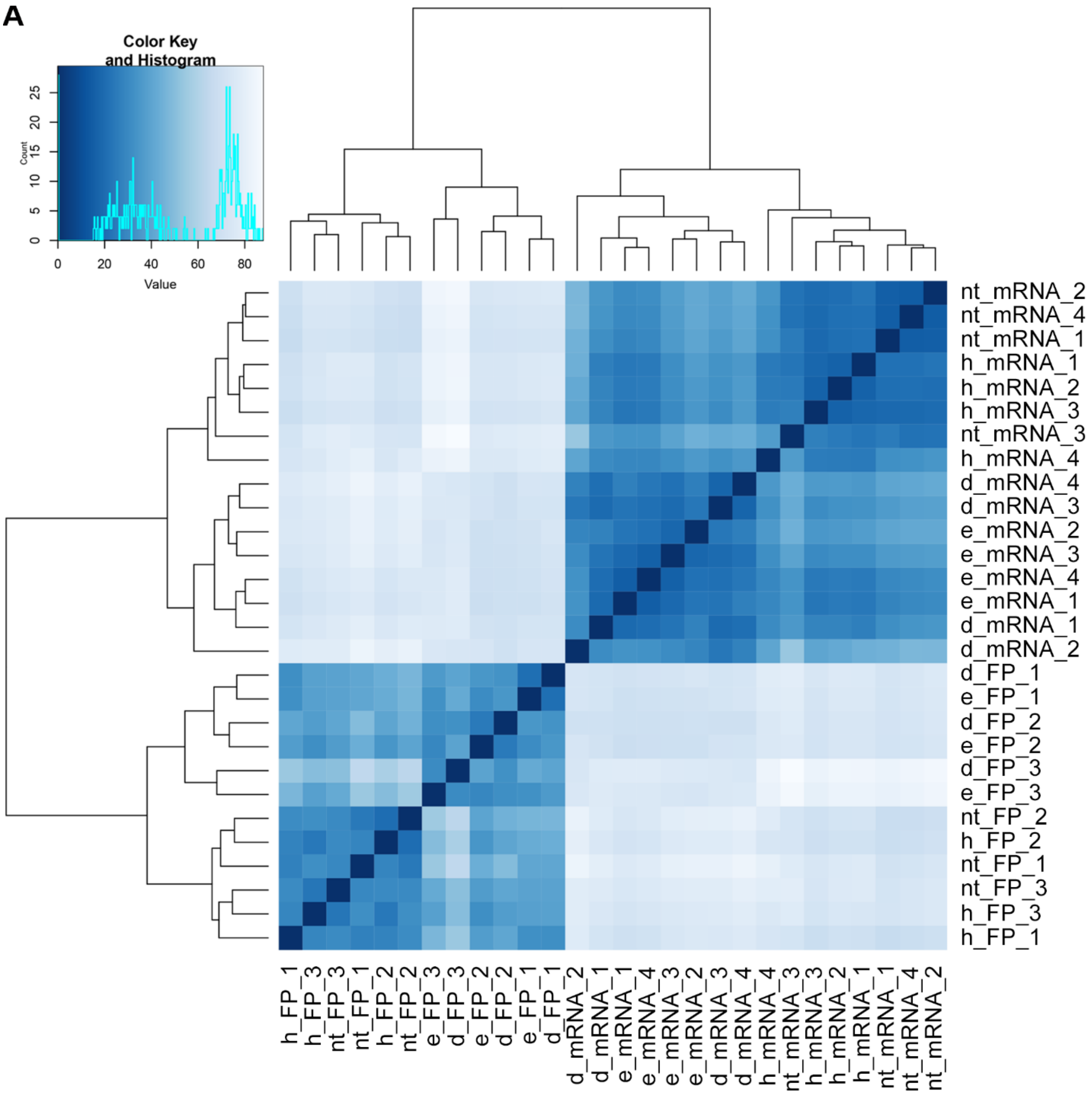


Supplementary Figure 4. **Clustering of Ribo-Seq and RNA-Seq libraries. (A)** Heatmap and dendrogram resulting from hierarchical clustering analysis of Ribo-Seq libraries and RNA-Seq libraries. Analysis was performed on gene counts by DESeq2 (Love *et al*, 2014) and biomaRt (Durinck *et al*, 2009).

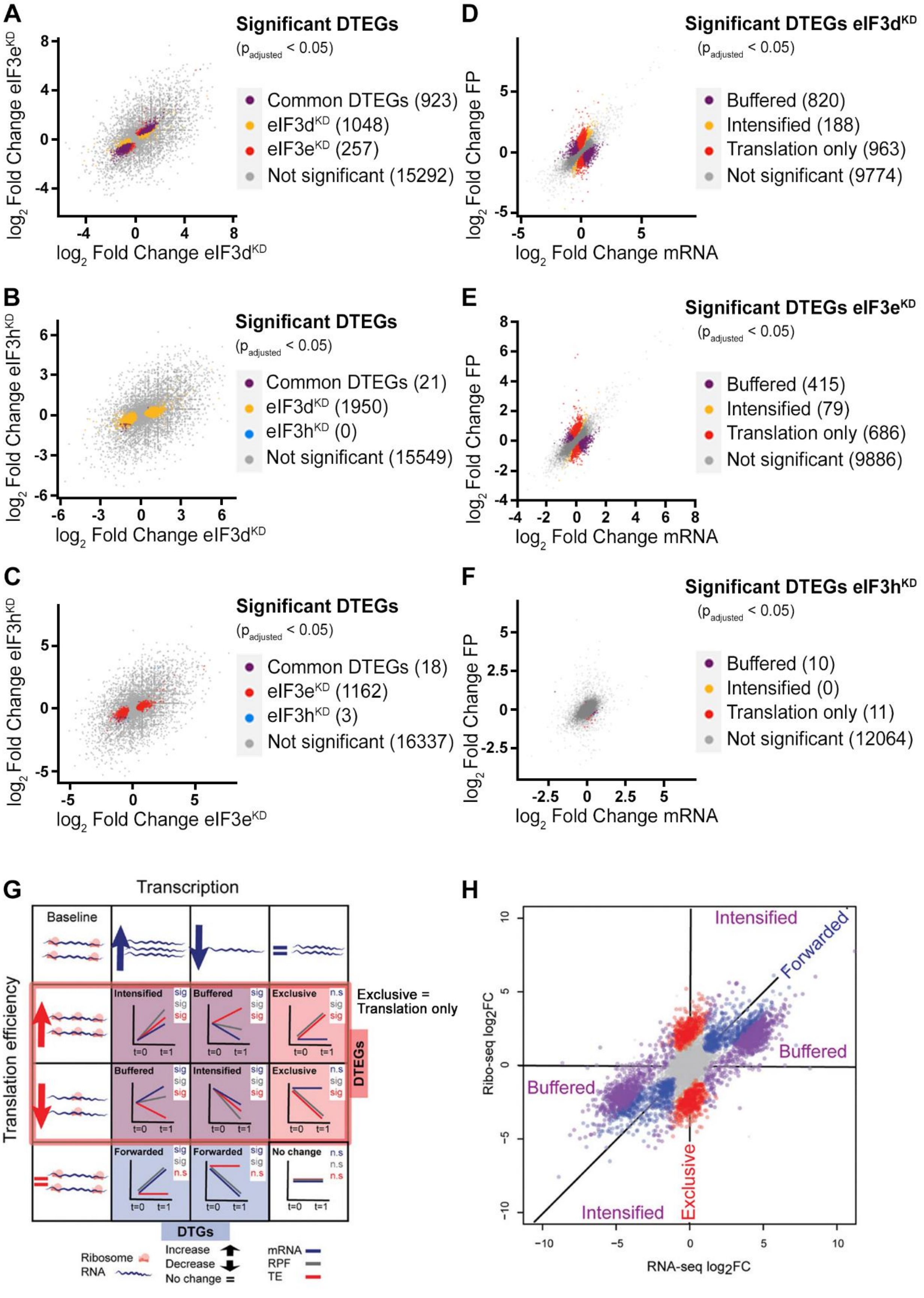
A
Significant DTEGs
(p adjusted < 0.05)
Common DTEGs (923)
eIF3d KD (1048)
eIF3e KD (257)
Not significant (15292)
log2 Fold Change eIF3e KD
log2 Fold Change eIF3d KD
B
Significant DTEGs
(p adjusted < 0.05)
Common DTEGs (21)
eIF3d KD (1950)
eIF3h KD (0)
Not significant (15549)
log2 Fold Change eIF3h KD
log2 Fold Change eIF3d KD
C
Significant DTEGs
(p adjusted < 0.05)
Common DTEGs (18)
eIF3e KD (1162)
eIF3h KD (3)
Not significant (16337)
log2 Fold Change eIF3h KD
log2 Fold Change eIF3e KD
D
Significant DTEGs eIF3d KD
(p adjusted < 0.05)
Buffered (820)
Intensified (188)
Translation only (963)
Not significant (9774)
log2 Fold Change FP
log2 Fold Change mRNA
E
Significant DTEGs eIF3e KD
(p adjusted < 0.05)
Buffered (415)
Intensified (79)
Translation only (686)
Not significant (9886)
log2 Fold Change FP
log2 Fold Change mRNA
F
Significant DTEGs eIF3h KD
(p adjusted < 0.05)
Buffered (10)
Intensified (0)
Translation only (11)
Not significant (12064)
log2 Fold Change FP
log2 Fold Change mRNA
G
Transcription
Baseline
Translation efficiency
Intensified
Buffered
Exclusive
Exclusive = Translation only
DTEGs
Forwarded
No change
DTGs
sig
n.s
t=0
t=1
Ribosome
RNA
Increase
Decrease
No change =
mRNA
RPF
TE
H
Intensified
Forwarded
Buffered
Exclusive
Ribo-seq log2FC
RNA-seq log2FC

Supplementary Figure 5. **DTEGs identified in all knock-downs largely overlap. (A-C)** Scatter plots of genes with significant TE changes and their overlap in individual knock-downs. **(A)** eIF3d$^{KD}$ vs. eIF3e$^{KD}$ **(B)** eIF3d$^{KD}$ vs. eIF3h$^{KD}$ **(C)** eIF3e$^{KD}$ vs. eIF3h$^{KD}$. **(D-F)** Scatter plots showing classification of significant DTEGs into different groups based on fold changes of FP and mRNA. Buffered DTEGs (purple) display a significant change in TE that counteracts the change in mRNA, hence buffering the effect of transcription. Translation only DTEGs (red) have a significant change in FP while maintaining the same mRNA levels, which results in a significant change in TE. Translationally intensified DTEGs (yellow) have a significant change in TE that occurs in the same direction as the effect of transcription (i.e., a gene exhibiting an increase in transcription also exhibits increase in TE that altogether boost protein production). **(D)** eIF3d$^{KD}$, **(E)** eIF3e$^{KD}$, **(F)** eIF3h$^{KD}$. (**G** - **H**) Classification of genes based on fold changes of FP (RFP), mRNA, and TE (adopted from (Chothani *et al*, 2019) with permission). (**G**) A gene could be either DTG (Differentially Transcribed Gene) and/or DTEG (Differential Translation Efficiency Gene), and based on the direction of change would fall into one of the eight gene-regulatory possibilities (sig: significant, n.s.: not significant). Translationally forwarded genes are DTGs that have a significant change in mRNA and FP at the same rate, with no significant change in TE. Conversely, translationally exclusive / translation only genes are DTEGs that have a significant change in FP, with no change in mRNA leading to a significant change in TE. Several genes are both DTGs and DTEGs, and their regulatory class is determined based on a combination of the relative direction of change between transcription and translation efficiency. Specifically, translationally buffered genes have a significant change in TE that counteracts the change in RNA; hence, buffering the effect of transcription. Translationally intensified genes have a significant change in TE that acts with the effect of transcription. In all cases, the change in RNA can be either positive or negative, and where buffering or intensifying takes place, the direction of change is taken into account. For example, a gene that exhibits an increase in transcription and an increase in translation efficiency is classified as intensified, while a gene that exhibits an increase in transcription but a decrease in translational efficiency is classified as buffered. (**H**) Simulated data showing fold changes for each gene in RNA-seq and Ribo-seq data. Translationally forwarded genes (in blue), exclusive (translation only) genes (in red), buffered genes (in purple), and intensified genes (in purple) are highlighted.

A

| KEGG term | # of genes in pathway | eIF3dKD all upregulated | | | eIF3eKD all upregulated | | | eIF3dKD/eIF3eKD common | | | eIF3dKD unique upregulated | | | eIF3dKD translation only upregulated | | | eIF3eKD translation only upregulated | | |
|---|---|---|---|---|---|---|---|---|---|---|---|---|---|---|---|---|---|---|---|
| | | rank | p-value | # of genes | rank | p-value | # of genes | rank | p-value | # of genes | rank | p-value | # of genes | rank | p-value | # of genes | rank | p-value | # of genes |
| Lysosome | 132 | 1 | 1.56E-34 * | 51 | 1 | 8.29E-22 * | 32 | 1 | 2.67E-25 * | 32 | 1 | 4.59E-10 * | 19 | 1 | 1.48E-26 * | 36 | 1 | 3.99E-18 * | 25 |
| Protein processing in ER | 171 | 2 | 1.13E-24 * | 48 | 2 | 6.21E-16 * | 30 | 2 | 4.64E-19 * | 30 | 2 | 3.05E-07 * | 18 | 2 | 5.94E-18 * | 32 | 2 | 5.33E-17 * | 27 |
| N-Glycan biosynthesis | 50 | 3 | 1.44E-20 * | 25 | 3 | 2.11E-13 * | 16 | 3 | 3.71E-15 * | 16 | 5 | 3.57E-06 * | 9 | 3 | 2.83E-12 * | 15 | 3 | 9.82E-12 * | 13 |
| Various types of N-glycan biosynthesis | 39 | 4 | 3.67E-13 * | 17 | 5 | 5.75E-09 * | 11 | 4 | 3.71E-10 * | 11 | 11 | 3.84E-04 * | 6 | 4 | 1.8E-11 * | 13 | 4 | 1.5E-10 * | 11 |
| ECM-receptor interaction | 88 | 5 | 8.19E-12 * | 23 | 7 | 9.18E-07 * | 13 | 6 | 4.61E-08 * | 13 | 9 | 6.87E-05 * | 10 | 6 | 7.81E-07 * | 13 | 6 | 1.69E-08 * | 13 |
| Other glycan degradation | 18 | 6 | 1.39E-09 * | 10 | 10 | 1.03E-04 * | 5 | 10 | 2.98E-05 * | 5 | 8 | 6.26E-05 * | 5 | 8 | 5.77E-06 * | 6 | 17 | 4.93E-03 | 3 |
| Protein digestion and absorption | 103 | 7 | 1.56E-09 * | 22 | 12 | 5.92E-04 * | 10 | 11 | 7.27E-05 * | 10 | 7 | 1.02E-05 * | 12 | 12 | 2.52E-05 * | 12 | 15 | 4.23E-03 * | 7 |
| Glycosaminoglycan degradation | 19 | 8 | 2.81E-09 * | 10 | 8 | 8.95E-06 * | 6 | 7 | 1.99E-06 * | 6 | 13 | 1.11E-03 * | 4 | 5 | 1.75E-08 * | 8 | 8 | 2.65E-05 * | 5 |
| GPI-anchor biosynthesis | 26 | 9 | 8.33E-09 * | 11 | 6 | 1.90E-08 * | 9 | 5 | 4.59E-08 * | 8 | 32 | 0.026359 | 3 | 7 | 4.67E-06 * | 7 | 7 | 9.14E-06 * | 6 |
| Glycosaminoglycan biosynthesis | 20 | 10 | 6.43E-07 * | 9 | 40 | NS | 3 | 34 | 0.026581 | 3 | 6 | 5.92E-06 * | 6 | 19 | 5.77E-04 * | 4 | 57 | NS | 1 |
| ABC transporters | 45 | 11 | 6.53E-07 * | 12 | 49 | NS | 3 | 43 | NS | 3 | 3 | 1.41E-06 * | 9 | 13 | 2.66E-05 * | 8 | 52 | NS | 2 |
| Phagosome | 152 | 13 | 2.13E-06 * | 22 | 11 | 3.33E-04 * | 13 | 9 | 2.44E-05 * | 13 | 27 | 0.014363 | 9 | 22 | 3.17E-03 * | 11 | 13 | 8.66E-04 * | 10 |
| Other types of O-glycan biosynthesis | 47 | 15 | 7.66E-06 * | 11 | 82 | NS | 2 | 70 | NS | 2 | 4 | 2.08E-06 * | 9 | 14 | 3.7E-05 * | 8 | 106 | NS | 1 |
| Cholesterol metabolism | 50 | 16 | 1.45E-05 * | 11 | 9 | 6.51E-05 * | 8 | 8 | 1.01E-05 * | 8 | 55 | NS | 3 | 10 | 7.5E-06 * | 9 | 10 | 4.24E-04 * | 6 |
| Protein export | 23 | 18 | 5.81E-05 * | 7 | 27 | 0.02503 | 3 | 46 | NS | 2 | 10 | 2.22E-04 * | 5 | 18 | 3.39E-04 * | 5 | 21 | 9.96E-03 | 3 |
| Bile secretion | 89 | 26 | 2.57E-04 * | 13 | 48 | NS | 5 | 37 | 0.04397 | 5 | 17 | 1.86E-03 * | 8 | 9 | 6.23E-06 * | 12 | 55 | NS | 3 |
| Coronavirus disease | 232 | 42 | 0.02348 | 18 | 16 | 2.21E-03 * | 15 | 55 | NS | 8 | 48 | NS | 10 | 35 | 0.026725 | 12 | 9 | 5.9E-05 * | 15 |
| Ribosome | 158 | 52 | NS | 11 | 4 | 5.2E-10 * | 22 | 25 | 7.07E-03 | 9 | 196 | NS | 2 | 46 | NS | 8 | 5 | 3.18E-10 * | 19 |

B

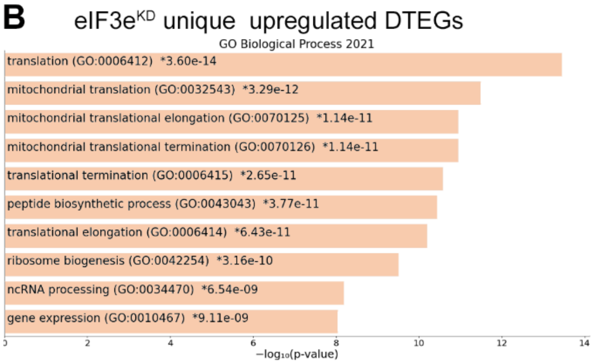


C

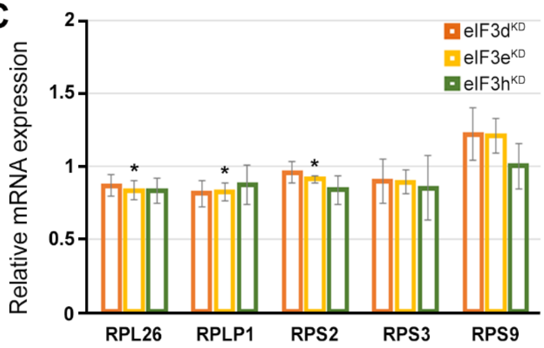


Supplementary Figure 6. **KEGG pathway enrichment analysis of the eIF3d$^{KD}$- and eIF3e$^{KD}$-associated upregulated DTEGs reveals upregulation of ribosomal proteins. (A)** List of top 10 significantly enriched KEGG pathways for eIF3d$^{KD}$ "all", eIF3e$^{KD}$ "all", eIF3d$^{KD}$/eIF3e$^{KD}$ "common", eIF3d$^{KD}$ "unique", eIF3d$^{KD}$ "translation only", and eIF3e$^{KD}$ "translation only" groups of upregulated DTEGs. Ranking corresponds to the lowest-to-highest p-value (calculated by Enrichr gene set search engine), with the "eIF3d$^{KD}$ all" group setting the primary ranking. Less significant terms ranking 11 and below are in grey. Terms specifically discussed in the main text are highlighted in green. For each term, the total number of genes in the pathway and number of DTEGs found in each pathway is indicated. Only significant p-values (< 0.05) are shown. NS = not significant p-value (> 0.05) Asterisk next to a p-value indicates that a given term also has a significant Benjamini-Hochberg adjusted p-value (< 0.05). **(B)** The bar chart shows the top 10 enriched GO Biological Process terms for eIF3e$^{KD}$ "unique upregulated" DTEGs, along with their corresponding p-values. **(C)** Relative mRNA expression of selected Ribosomal protein genes normalized to NT control = 1. One-sample t-test was used for statistical evaluation, * = P<0.05.

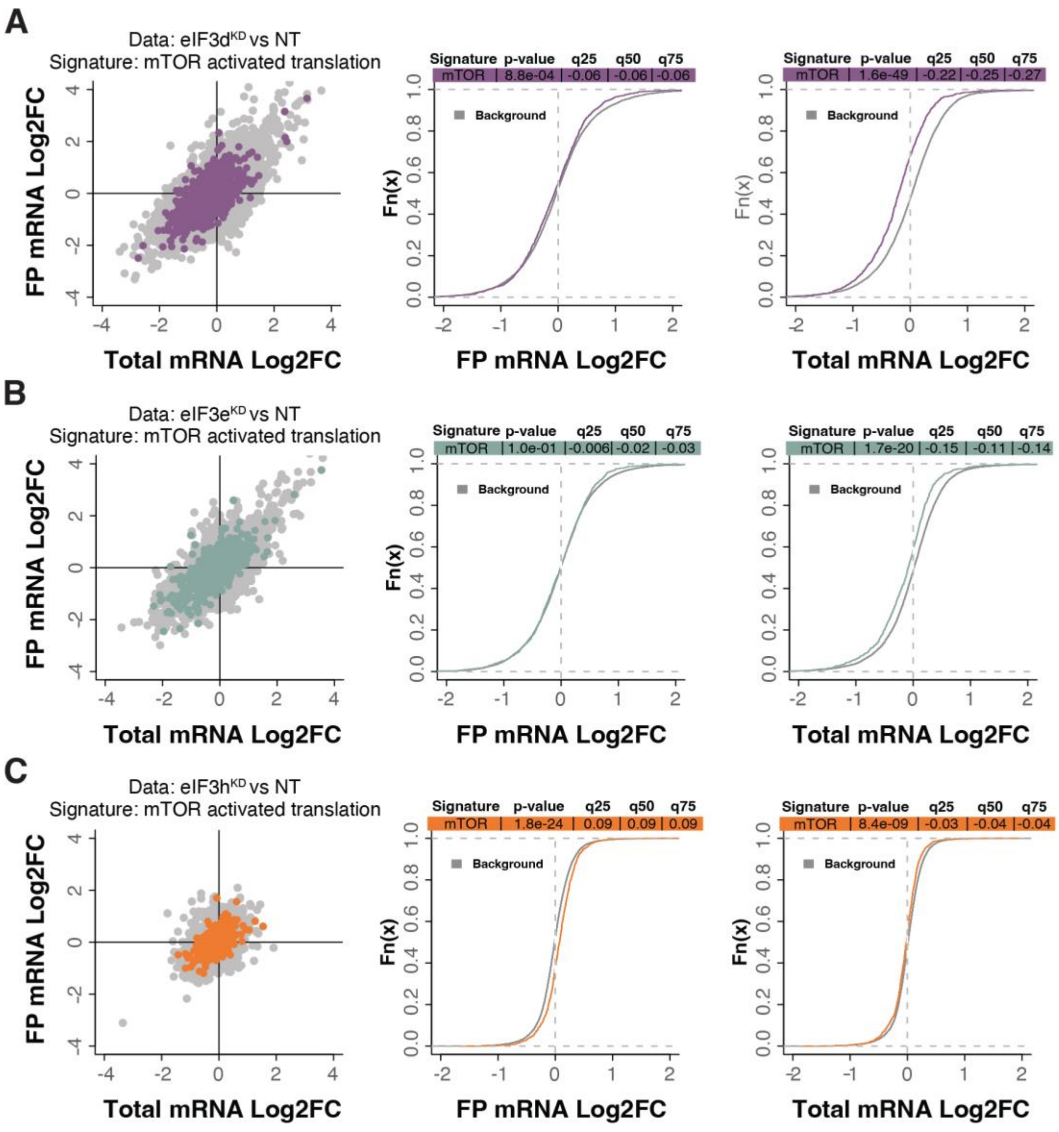


Supplementary Figure 7. **mTOR-sensitive transcripts tend to be translationally offset (buffered) with loss of eIF3d and eIF3e. (A-C)** Scatterplots from translatome analysis of eIF3d$^{KD}$ **(A)**, eIF3e$^{KD}$ **(B)**, and eIF3h$^{KD}$ **(C)** with the location of transcripts that are translationally activated by insulin stimulation (mTOR activation) (Gandin *et al*, 2016) colored (left panels). Middle and right panels show the empirical cumulative distribution functions of $\log_2$ fold changes in FP and total mRNA for the transcripts that are translationally activated by mTOR signaling. The background constituting all other transcripts are shown as grey curves. Significant differences between the distributions were identified using the Wilcoxon rank-sum test. Differences between the distributions at each quantile are indicated. mRNAs of mTOR-sensitive transcripts in eIF3d$^{KD}$ and eIF3e$^{KD}$ display significant downregulation (shift to the left) while the FPs remain unchanged, suggesting translational buffering.

**A**

| KEGG term | # of genes in pathway | eIF3dKD all downregulated rank | eIF3dKD all downregulated p-value | eIF3dKD all downregulated # of genes | eIF3eKD all downregulated rank | eIF3eKD all downregulated p-value | eIF3eKD all downregulated # of genes | eIF3dKD/eIF3eKD common downregulated rank | eIF3dKD/eIF3eKD common downregulated p-value | eIF3dKD/eIF3eKD common downregulated # of genes |
|---|---|---|---|---|---|---|---|---|---|---|
| Chronic myeloid leukemia | 76 | 1 | 1.652E-10 | 21 | 10 | 7.793E-08 | 14 | 7 | 4.321E-09 | 14 |
| Neurotrophin signaling pathway | 119 | 2 | 3.414E-10 | 26 | 2 | 4.614E-09 | 19 | 9 | 5.603E-09 | 17 |
| Autophagy | 141 | 3 | 3.734E-10 | 29 | 1 | 1.538E-09 | 21 | 2 | 1.183E-09 | 19 |
| MAPK signaling pathway | 294 | 4 | 2.572E-09 | 42 | 5 | 1.221E-08 | 30 | 12 | 2.115E-08 | 26 |
| Yersinia infection | 137 | 5 | 8.208E-09 | 26 | 4 | 8.945E-09 | 20 | 3 | 1.183E-09 | 19 |
| Oocyte meiosis | 131 | 6 | 1.017E-08 | 25 | 11 | 1.004E-07 | 18 | 11 | 1.941E-08 | 17 |
| Ubiquitin mediated proteolysis | 142 | 7 | 1.316E-08 | 26 | 9 | 7.000E-08 | 19 | 10 | 1.103E-08 | 18 |
| Dopaminergic synapse | 132 | 8 | 1.654E-08 | 25 | 3 | 4.637E-09 | 20 | 1 | 6.206E-10 | 19 |
| Fc gamma R-mediated phagocytosis | 97 | 9 | 2.002E-08 | 21 | 7 | 4.742E-08 | 16 | 4 | 1.845E-09 | 16 |
| Renal cell carcinoma | 69 | 10 | 5.914E-08 | 17 | 14 | 1.722E-07 | 13 | 18 | 1.085E-07 | 12 |
| Human cytomegalovirus infection | 225 | 11 | 6.852E-08 | 33 | 6 | 3.805E-08 | 25 | 5 | 1.899E-09 | 24 |
| T cell receptor signaling pathway | 104 | 14 | 3.406E-07 | 20 | 13 | 1.298E-07 | 16 | 8 | 5.269E-09 | 16 |
| Shigellosis | 247 | 20 | 1.661E-06 | 33 | 8 | 5.666E-08 | 26 | 6 | 2.391E-09 | 25 |

**B**

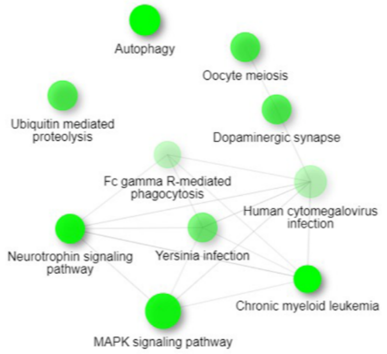


**C**

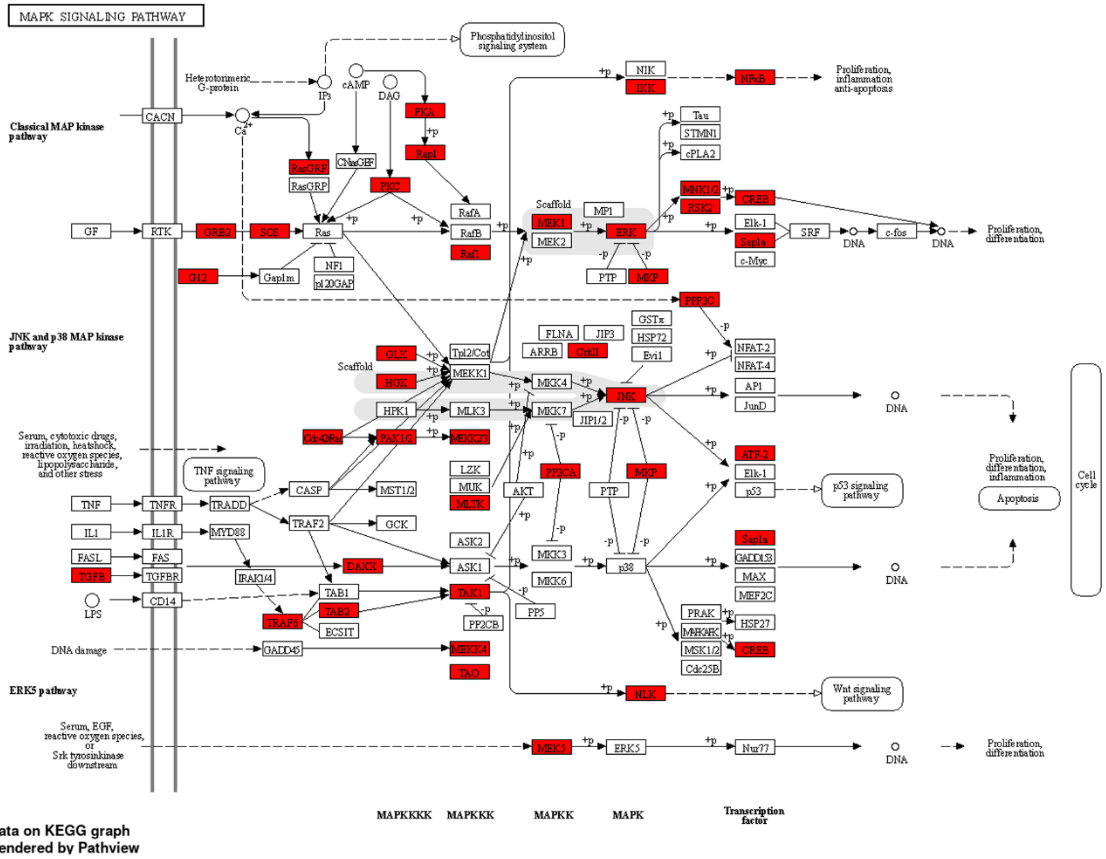

Supplementary Figure 8. **KEGG pathway enrichment analysis of the eIF3d$^{KD}$- and eIF3e$^{KD}$-associated downregulated DTEGs reveals downregulation of “MAPK signaling pathway” components. (A)** List of top 10 significantly enriched KEGG pathways for eIF3d$^{KD}$ “all”, eIF3e$^{KD}$ “all”, and eIF3d$^{KD}$/eIF3e$^{KD}$ “common” groups of downregulated DTEGs. Ranking corresponds to the lowest-to-highest p-value (calculated by Enrichr gene set search engine), with the eIF3d$^{KD}$ “all” group setting the primary ranking. Less significant terms ranking 11 and below are in grey. For each term, the total number of genes in the pathway and number of DTEGs found in each pathway is indicated. All results presented have also significant adjusted p-values as calculated using the Benjamini-Hochberg (BH) procedure to account for multiple hypotheses. **(B)** Network graph of the top 10 most significantly enriched KEGG terms for the eIF3d$^{KD}$ “all downregulated” group of DTEGs. Related KEGG terms are connected with a solid line if they share 20% or more genes. Darker nodes represent more significantly enriched gene sets. Greater nodes represent larger gene sets. **(C)** The KEGG MAPK signaling pathway hsa04010 scheme. Downregulated DTEGs in eIF3d$^{KD}$ matching the genes of this pathway are highlighted in red. See text and Supplementary Spreadsheet S2 for further details.

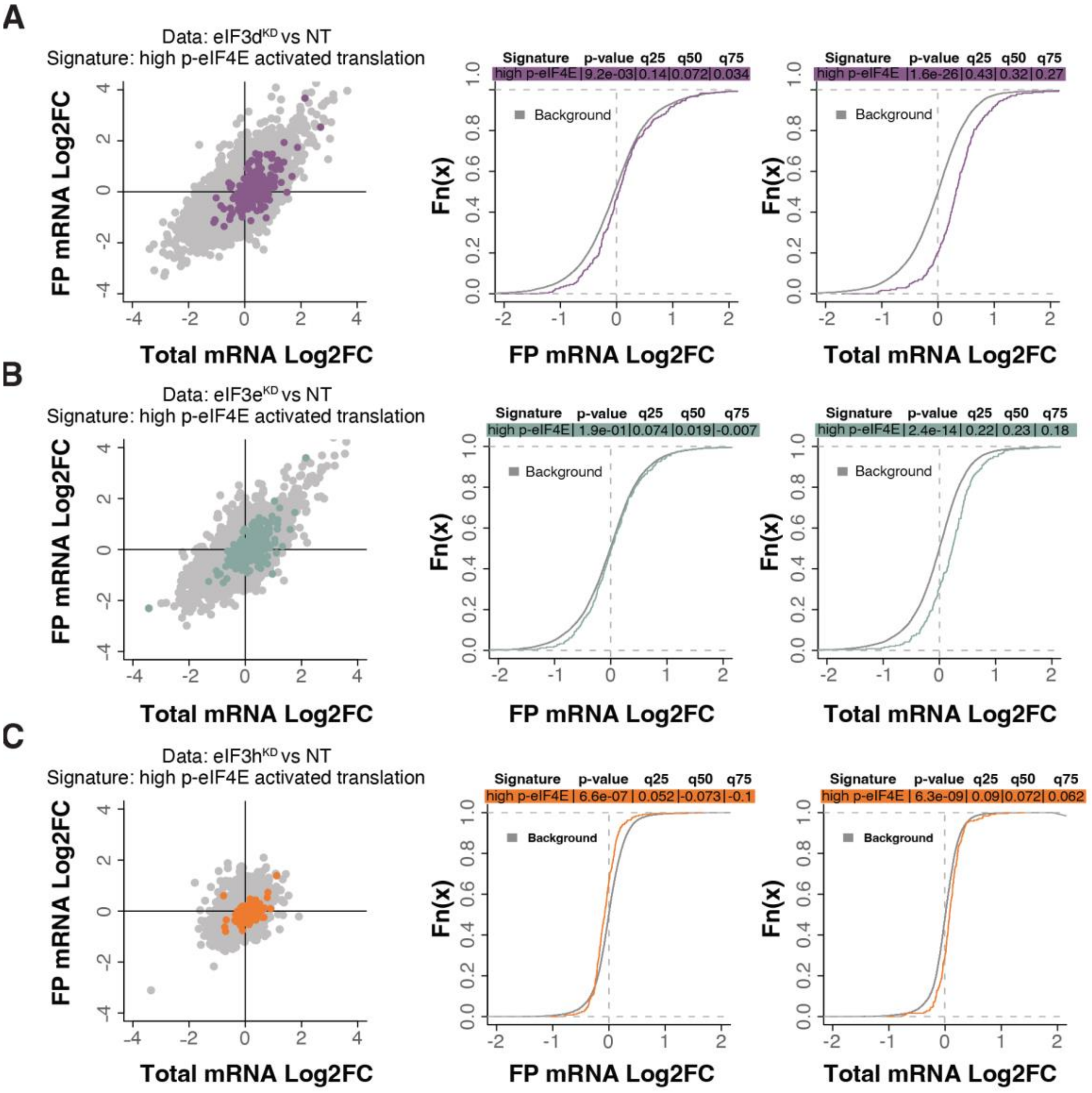


Supplementary Figure 9. **Loss of eIF3 subunits leads to translational offsetting of transcripts with enhanced translation downstream of phosphorylated eIF4E. (A-C)** Scatterplots from translatome analysis of eIF3d$^{KD}$ **(A)**, eIF3e$^{KD}$ **(B)**, and eIF3h$^{KD}$ **(C)** with the location of transcripts translationally activated by high phosphorylation of eIF4E (Karampelias *et al*, 2022) colored (left panels). Middle and right panels show the empirical cumulative distribution functions of $\log_2$ fold changes in FP and total mRNA for the p-eIF4E sensitive transcripts. The background constituting all other transcripts are shown as grey curves. Significant differences between the distributions were identified using the Wilcoxon rank-sum test. Differences between the distributions at each quantile are indicated. mRNAs of phospho-eIF4E sensitive transcripts in eIF3d$^{KD}$ and eIF3e$^{KD}$ display significant upregulation (shift to the right) while the FPs remain unchanged, suggesting translational buffering.

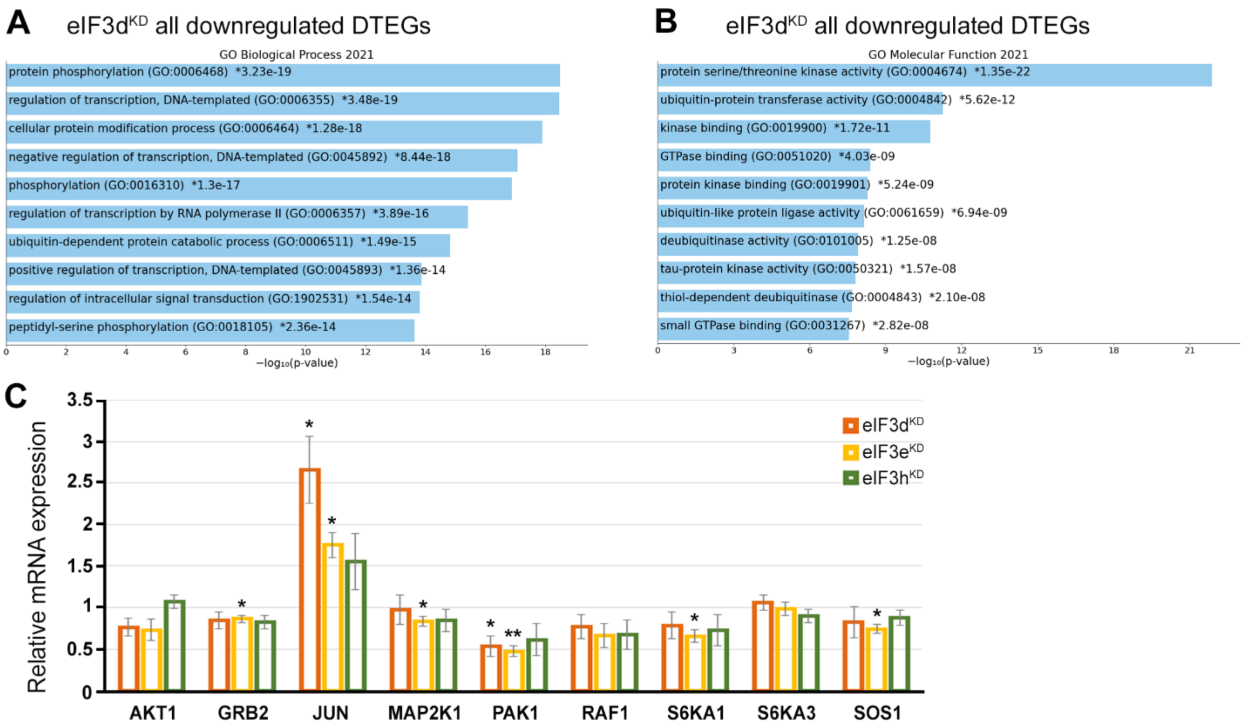


Supplementary Figure 10. **GO enrichment analysis for eIF3d$^{KD}$ associated downregulated DTEGs. (A)** The bar chart shows the top 10 enriched GO Biological Process terms for eIF3d$^{KD}$ “all downregulated” DTEGs, along with their corresponding p-values. **(B)** The bar chart shows the top 10 enriched GO Molecular Function terms for eIF3d$^{KD}$ “all downregulated” DTEGs, along with their corresponding p-values. **(A-B)** Colored bars correspond to terms with significant p-values (<0.05). Asterisk indicates that a given term also has a significant adjusted p-value (<0.05). All P-values were calculated by Enrichr gene set search engine. **(C)** Relative mRNA levels of selected genes from MAPK/ERK signaling pathway normalized to NT control = 1. One-sample t-test was used for statistical evaluation, P-values: * = P<0.05, ** = P<0.01.

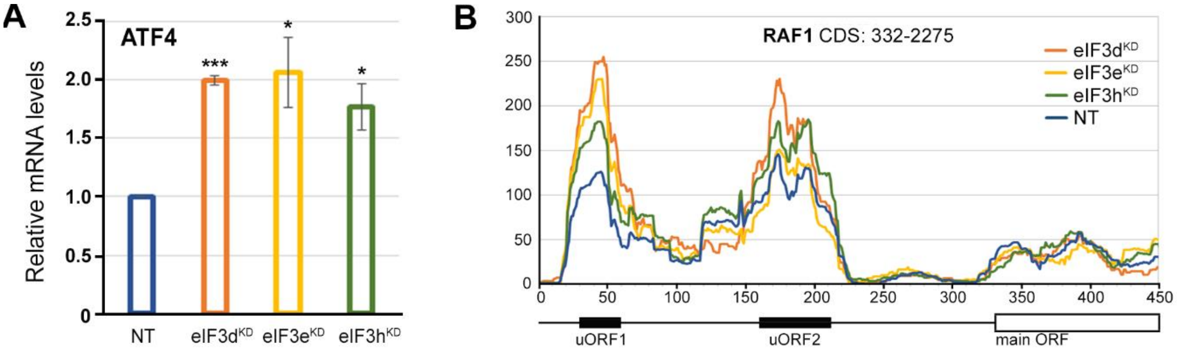


Supplementary Figure 11. **The *ATF4* mRNA is upregulated in all three eIF3 knock-downs tested. (A)** Relative mRNA levels of ATF4 transcription factor. One-sample t-test was used for statistical evaluation, * = P<0.05, *** = P<0.001. **(B)** Normalized ribosomal footprint coverage along the *RAF1* mRNA (first 450 nucleotides). Schematic with regulatory elements is shown at the bottom. Average of all three replicates is shown. Footprint coverage was normalized to all footprints mapping to *RAF1* mRNA.

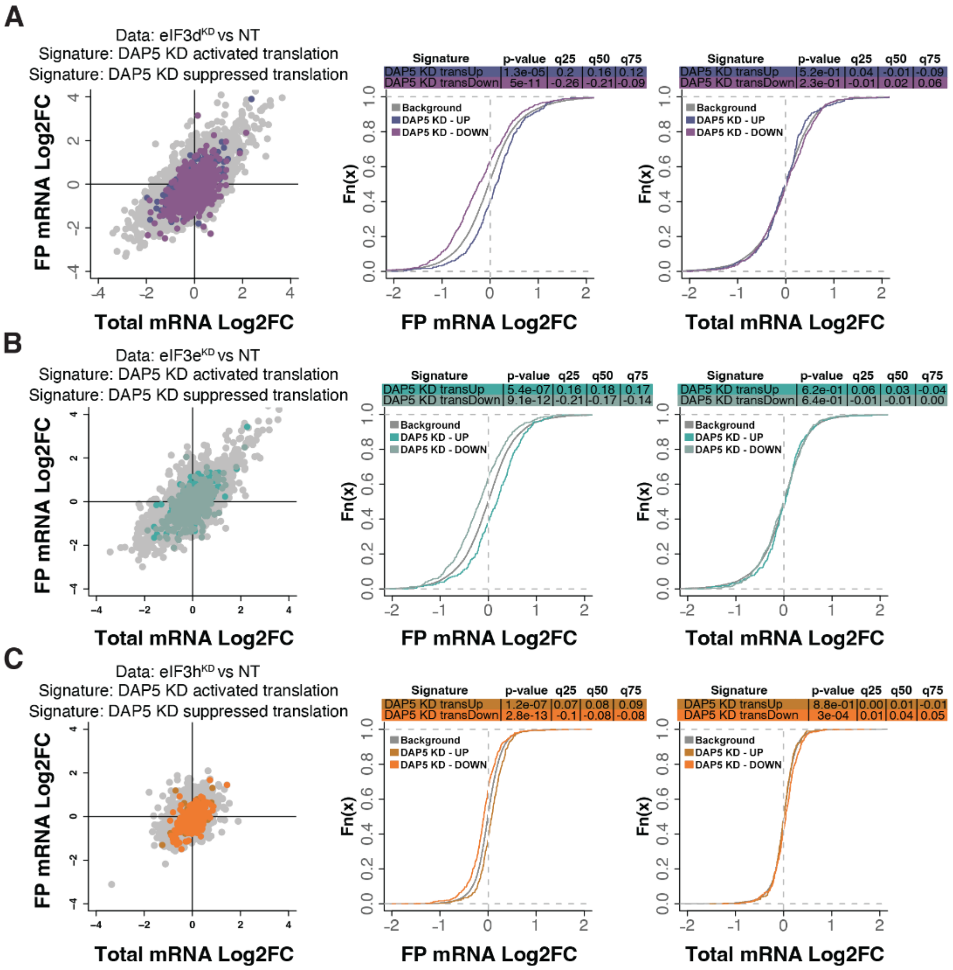


Supplementary Figure 12. **Loss of eIF3 subunits modulates signatures of DAP5-dependent translation. (A-C)** Scatterplots from translatome analysis of eIF3d$^{KD}$ **(A)**, eIF3e$^{KD}$ **(B)**, and eIF3h$^{KD}$ **(C)** with the location of transcripts that are translationally activated and suppressed by KD of DAP5 (David *et al*, 2022) colored (left panels). Middle and right panels show the empirical cumulative distribution functions of $\log_2$ fold changes in FP and total mRNA for the transcripts that are translationally activated or suppressed upon loss of DAP5. The background constituting all other transcripts are shown as grey curves. Significant differences between the distributions were identified using the Wilcoxon rank-sum test. Differences between the distributions at each quantile are indicated.

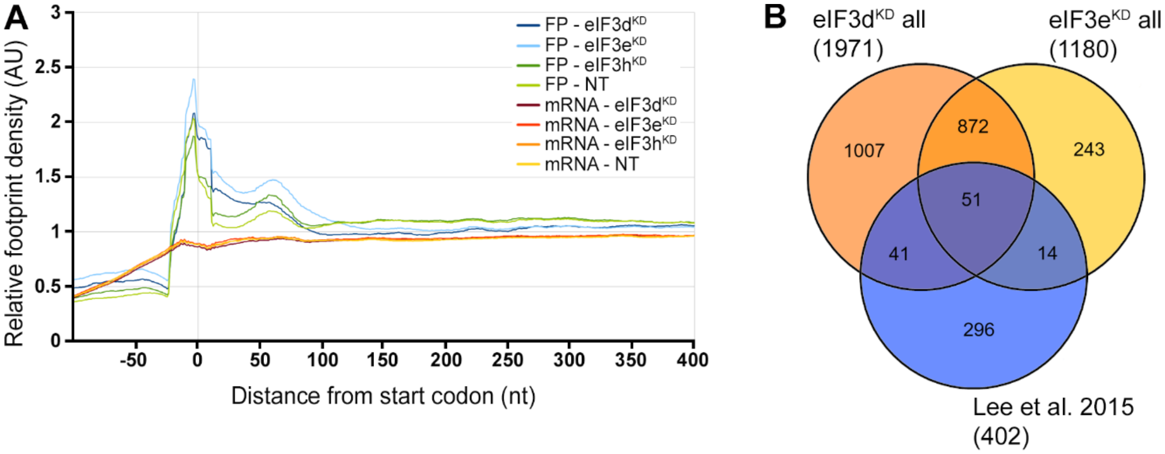


Supplementary Figure 13. **eIF3d$^{KD}$, eIF3e$^{KD}$, and eIF3h$^{KD}$ do not show accumulation of footprints at the beginning of CDSes, as reported previously, but display a significant overlap with mRNAs directly interacting with eIF3. (A)** Metagene plot of ribosome density distribution around the start codon of all mRNAs in eIF3d$^{KD}$, eIF3e$^{KD}$ and eIF3h$^{KD}$. The plot shows averages of triplicates from Ribo-Seq and quadruplicates from RNA-Seq. **(B)** Venn diagram of the overlap between eIF3d/e$^{KD}$ DTEGs and mRNAs directly interacting with eIF3 as determined by the CLIP assay in (Lee *et al*, 2015).

**Supplementary Table S1.** siRNAs used in the study.

| ON-TARGETplus siRNA DHARMACON | cat # |
|---|---|
| eIF3d | L-017556-00 |
| eIF3e | L-010518-00 |
| eIF3h | L-003883-00 |
| Non-targeting | D-001810-03 |

**Supplementary Table S2.** qPCR primers used in the study.

| Name | 5ˈto 3ˈ sequence |
|---|---|
| Akt1* | CTTCTTTGCCGGTATCGTG |
| Akt1-R* | TGCTGTCATCTTGGTCAGG |
| ALAS1 | CCACTGGAAGAGCTGTGTGATGTG |
| ALAS1-R | GCGATGTACCCTCCAACACAACC |
| ATF4 | AAACCTTACGATCCTCCTGGAG |
| ATF4-R | TGGCTGCTGTCTTGTTTTGC |
| eIF3b* | TGTGAAAGGTACCTGGTGAC |
| eIF3b R* | AATAGGCCAATGGGCTGAG |
| eIF3d | CCAACCCAAACCCGTTTGTG |
| eIF3d R | TCTTCAGCTCCGTGGCAATG |
| eIF3e | CTGGTTCCAGCAACAGATAG |
| eIF3e R | GTGGCTGATAAAGGAAGAGG |
| eIF3h* | GCTGACTTTGATGAAGTCCA |
| eIF3h R* | ATGTGGACTGATACCAGCC |
| GRB2* | CTCTGTCAAGTTTGGAAACGA |
| GRB-R* | TGAACTTCACCACCCAGAG |
| Jun* | CAACATGCTCAGGGAACAG |
| Jun-R* | ACTGTTAACGTGGTTCATGAC |
| MAP2K1* | CCAGAAAGCTAATTCATCTGGAG |
| MAP2K1-R* | GTTGCACTCATGCAGAACC |
| PAK1* | CTTTGACCCGGAATACTGAG |
| PAK1-R* | CACTATGCTTCGTAATTTCTCC |
| RAF1* | CACAACTTTGCTCGGAAGAC |
| RAF1-R* | ACATCGAAATCCATTGAGCAG |
| RPL26* | GATGATGAAGTTCAGGTTGTACG |
| RPL26-R* | CATATTTCTTCCTGTAAACCTGGAC |
| RPLP1* | CTACTCGGCCCTCATTCTG |
| RPLP1-R* | CCACTTTCTTCTCCTCAGC |
| RPS2* | AGATCATTGATTTCTTCCTGGG |

| | |
|---|---|
| RPS2-R* | CAACAAATGCCTTGAACCTG |
| RPS3* | TTATCTTAGCCACCAGAACAC |
| RPS3-R* | TCTTCTGAACTACAGCAGTC |
| RPS6KA1* | AACGCTGAAAGTACGTGAC |
| RPS6KA1-R | CATAGTGCAGCTTCACCAC |
| RPS6KA3* | CTATACAATGCTTACCGGTTACAC |
| RPS6KA3-R* | CTACCTATTCGTGCCAATATTTCC |
| RPS9* | AAGAGCTGAAGCTGATCGG |
| RPS9-R* | AATTTGACCCTCCAGACCTC |
| SOS1* | TGGTGTCCTTGAGGTTGTC |
| SOS1-R* | GGCGACTTGGTATTTGCTC |

*primersˈ sequences were obtained from the database GETPrime, available at http://bbcftools.epfl.ch/getprime

**Supplementary Table S3.** Antibodies used in the study.

| Antibody | source |
|---|---|
| AKT1 | Cell Signaling # 2938 |
| ATF4 | Cell Signaling # 11815 |
| eIF3b | Thermo Scientific # PA5-23278 |
| eIF3d | Sigma # HPA066216 |
| eIF3e | Thermo Scientific # PA5-29487 |
| eIF3h | Cell Signalling # 3413 |
| ERK1/2 | Santa Cruz # sc-514302 |
| P-ERK1/2 | Cell Signaling # 4370 |
| GAPDH | ThermoScientific # PA1-987 |
| GRB2 | BD Transduction Laboratories # 610112 |
| JUN | Cell Signaling # 9165 |
| P-JUN | Cell Signaling # 9261 |
| LAMIN B1 | Cell Signaling #12586 |
| MDM2 | Cell Signaling # 86934 |
| MEK1 | BD Transduction Laboratories # 610122 |
| PAK1 | Santa Cruz # sc-881 |
| RAF | Santa Cruz # sc-133 |
| RPL13A | Cell Signaling # 2765 |
| RPL26 | Cell Signaling # 5400 |
| RPLP1 | Sigma # HPA003368 |
| RPS2 | Santa Cruz # sc-130399 |
| RPS3 | Proteintech # 15198-1-AP |
| RPS9 | Thermo Scientific # PA5-13569 |

| | |
|---|---|
| RSK1 | Santa Cruz # sc-231 |
| RSK2 | Santa Cruz # sc-9986 |
| SOS1 | Cell Signaling # 12409 |

**Supplementary Table S4.** eIF3h$^{KD}$ "all downregulated" DTEGs, top 8 significant results from KEGG pathways as evaluated by Enrichr.

| Term | p-value | Overlap genes |
|---|---|---|
| Prostate cancer | 0.003773 | MDM2, ATF4 |
| Aldosterone synthesis and secretion | 0.003849 | PRKD3, ATF4 |
| Viral carcinogenesis | 0.015649 | MDM2, ATF4 |
| Human cytomegalovirus infection | 0.018997 | MDM2, ATF4 |
| Endocytosis | 0.023483 | MDM2, RAB11FIP2 |
| Bladder cancer | 0.038256 | MDM2 |
| PI3K-Akt signaling pathway | 0.043814 | MDM2, ATF4 |
| Cocaine addiction | 0.045558 | ATF4 |

**Supplementary Table S5.** eIF3h$^{KD}$ "all downregulated" DTEGs, top 10 significant results from GO Biological Process as evaluated by Enrichr.

| Term | p-value | Overlap genes |
|---|---|---|
| Protein ubiquitination | 0.001329 | RNF10, RMND5A, WAC, MDM2 |
| Mitotic G1 DNA damage checkpoint signaling | 0.001716 | WAC, MDM2 |
| Protein autoubiquitination | 0.001716 | RNF10, MDM2 |
| Protein deubiquitination | 0.001946 | ZRANB1, USP38, MDM2 |
| Protein modification by small protein removal | 0.002139 | ZRANB1, USP38, MDM2 |
| Proteolysis involved in cellular protein catabolic process | 0.002217 | ZRANB1, MDM2 |
| RNA splicing | 0.003849 | IVNS1ABP, SNRPG |
| Regulation of proteasomal ubiquitin-dependent protein catabolic process | 0.004162 | WAC, MDM2 |
| Ubiquitin-dependent protein catabolic process | 0.004316 | ZRANB1, RMND5A, MDM2 |
| Protein K29-linked deubiquitination | 0.004741 | ZRANB1 |
